# Investigating the use of odour and colour foraging cues by rosy-faced lovebirds (*Agapornis roseicollis*) using deep-learning based behavioural analysis

**DOI:** 10.1101/2024.02.18.580921

**Authors:** Winson King Wai Tsang, Emily Shui Kei Poon, Chris Newman, Christina D. Buesching, Simon Yung Wa Sin

## Abstract

Olfaction and vision can play important roles in optimizing foraging decisions of birds, enabling them to maximize their net rate of energy intake while searching for, handling, and consuming food. Parrots have been used extensively in avian cognition research, and some species use olfactory cues to find food. Here we pioneered machine learning analysis and pose-estimation with convolutional neural networks (CNNs) to elucidate the relative importance of visual and olfactory cues for informing foraging decisions in the rosy-faced lovebird (*Agapornis roseicollis*) as a non-typical model species. In a binary choice experiment, we used markerless body pose tracking to analyse bird response behaviours. Rosy-faced lovebirds quickly learnt to discriminate the feeder provisioned with food by forming an association with visual (red/green papers) but not olfactory (banana/almond odour) cues. When visual cues indicated the provisioned and empty feeders, feeder choice was more successful, choice latency shorter, and interest in the empty feeder significantly lower. This demonstrates that visual cues alone are sufficient to inform lovebird foraging decisions without needing to use olfactory cues, suggesting that selection has not driven olfactory-based foraging in lovebird evolution.

## Introduction

Optimal foraging theory posits that animals seek to maximize their net rate of energy intake while searching for, handling, consuming, and digesting food (Stephens & Krebs, 2019). Foraging optimality therefore depends not only on extrinsic variables, such as food availability, patch size, predator avoidance, and environmental stochasticity, but also on the forager’s ability to detect food (Martin, 2020). This may involve multiple sensory systems (i.e., detecting visual, auditory, tactile, and olfactory cues) integrated by cognitive processes (Talsma et al., 2010). Modalities may synergize each other, or one modality may have primacy. Depending on the specific conditions, feeding behaviour can thus be moderated by top-down factors, such as previous experience and the goals and expectations of the receiver, and by bottom-up factors, such as signal salience and detection threshold (Sumner and Sumner, 2020). When cognitive processing related to a specific task is more efficient in one modality than in another (e.g., when one modality is masked by environmental noise), the principle of ‘modality appropriateness’ applies (Welch and Warren, 1980). Ultimately, better understanding how animals engage their senses during ecologically important tasks may therefore inform on potential for adaptation to environmental change, where the growing appreciation of the role played by sensory abilities is driving a paradigm shift in foraging ecology (LaScala-Gruenewald et al., 2019).

The role of visual perception in avian foraging decisions is relatively well understood (Fernández-Juricic et al., 2004). Birds perceive wavelengths from 300 to 700 nm (Toomey et al., 2016), extending into the ultraviolet spectrum (Jacobs, 1992). Birds have four cone cell types and see more hues than humans (Toomey et al., 2016). This superior visual perception may support complex decision making, including foraging (e.g., Shrestha et al., 2013). Acoustic perception is also well-understood across bird taxa, although, outside of scavenging (Jackson et al., 2020), audition appears less important to foraging decisions (Elie et al., 2020).

In contrast, far less is known about avian olfaction, although birds use odour cues in individual discrimination (predators, relatives, partners, offspring, hetero-/conspecifics; Caro et al., 2015), nest recognition (Krause & Caspers, 2012), sexual advertisement (Caro et al., 2015), homing and navigation (Thorup et al., 2007), and foraging (e.g., Mäntylä et al., 2020; Rubene et al., 2019). For instance, insectivorous birds use herbivore-induced plant volatiles (HIPVs) in combination with visual cues to identify insect-damaged trees (Amo et al., 2013), and various bird species use both vision and olfaction either hierarchically or in combination to identify foraging sites (Rubene et al. 2019), while great tits (*Parus major*) can identify herbivore-damaged trees without any arthropod prey cues by using olfaction alone, but not vision alone (Amo et al., 2013). According to the dispersal syndrome hypothesis, fruits have evolved specific traits to attract dispersers (Lei et al. 2021), including fruit colour that signals higher lipid content and appeals particularly to avian dispersers. This suggests that birds rely more on visual cues than on odour or taste to detect food, although this may be influenced by species-specific fruit consumption techniques (Levey, 1987). Additionally, the olfactory receptor (OR) subgenome and its expression vary with olfactory ability and are shaped by ecological factors and life-history adaptations (Sin et al., 2022; Steiger et al. 2010).

Parrots (Psittaciformes) have been used extensively in avian cognition research (see Auersperg and von Bayern, 2019). Nevertheless, despite relying predominantly on fruits, seeds, and nectar (Ndithia & Perrin, 2009a; Toda et al., 2021), their ability to detect these foods using olfaction alone or in a multimodal combination with visual cues remains largely untested. Earlier studies (Healy and Guilford, 1990) used olfactory bulb ratio as an indicator of olfactory ability (Corfield et al., 2015), and inferred that parrots likely have a poor sense of smell. Nevertheless, studies showing that Yellow-backed chattering lories (*Lorius garrulus flavopallia*) (Roper, 2003), kakapo (*Strigops habroptilus*) (Gsell, 2012), kea (*Nestor notabilis*), and New Zealand kākā (*Nestor meridionalis*) (Gsell et al., 2016) use olfactory cues to find food have refuted this assumption.

To elucidate the role of olfaction and any multimodality between olfaction and vision in parrots, we conducted a food reward experiment testing the ability of rosy-faced lovebirds (*Agapornis roseicollis*), a common pet parrot species (Chan et al. 2020), to associate odour and colour cues with food presence/absence. Specifically, we investigated whether *A. roseicollis* can locate food rewards purely from i) olfactory cues, ii) visual cues, and iii) whether their success rate is enhanced if visual and olfactory cues are presented in a co-modal combination. To analyse decision-making processes in detail, we applied a machine learning analytical approach, using DeepLabCut (Mathis et al. 2018) and Simple Behavioural Analysis (SimBA) software (Nilsson et al., 2020), able to detect markerless posture estimation networks (convolutional neural networks) to classify behaviour from video, frame-by-frame. This approach can test nonlinear dependencies and unknown interactions across multiple variables unencumbered by the inductive bias implicit in a priori hypothesis testing (Sturman et al. 2020). We discuss our results in the context of optimal foraging theory to extend understanding of the ecological implications of avian cognition and sensory systems, where evidence that either of these sensory modalities, separately or in co-modality, enhance net energy gain or reduce the time taken to achieve that gain would support optimal foraging.

## Materials and methods

### Study species

We used 26 captive, sexually mature *A. roseicollis* (17 males, 9 females), kept at the Centre for Comparative Medicine Research (CCMR) animal facility at the University of Hong Kong. Birds were provisioned with artificial food pellets (Mazuri Small Bird Maintenance Diet 56A6), which did not contain any fruit and/or nut ingredients. Experiments were conducted in individual cages (60cm × 40cm × 35cm) under 6500K illumination (LED T5 tube, 7W, SUNSHINE).

### Experimental design and data collection

We conducted binary choice experiments, commonly used to study sensory discrimination in birds (e.g., Potier et al., 2021; Abankwah et al., 2020). Because lovebird natural diet includes fruits and seeds (Ndithia & Perrin, 2009a), we used natural sunflower seeds, Parakeet Higgins Vita Seed, and Mazuri food pellets as food reward in the experimental set-up.

To test sensory preferences, we designed a foraging task where birds had to select between two cylindrical (diameter × height: 5cm × 5cm), non-transparent feeders attached to a perch (20 cm) that allowed the bird to move freely between feeders and select between foraging cues (Supplementary Figure 1A). One feeder contained food (‘feeder_food_’), the other feeder was empty (‘feeder_w/o food_’). Both feeder bowls were completely covered with paper folded around the rim and fixed with a cable tie, preventing birds seeing the food reward. Red paper signified the feeder_food_, green paper signified the feeder_w/o food_, while white paper could cover both reward and empty feeders. A scent stick was attached to these paper cover, either untreated or treated with two drops of an odour cue: Banana scent (RAYNER’S) to indicate the feeder_food_, or 2 drops of almond scent (RAYNER’S) to indicate the feeder_w/o food_. These scents were chosen based on their successful application in similar studies.

Almond odour occurs naturally, associated with toxicity in plants, and has been used successfully in avoidance learning experiments in chickens (*Gallus gallus domesticus*) (Roper and Marples, 1997). Banana odour was used because toucans (*Ramphastos* spp.), scarlet macaws (*Ara macao)* (Hernández et al. 2022), and red-winged starlings (*Onychognathus morio*) (Zungu et al. 2014) can detect and use it when making foraging decisions (see Supplementary Material for habituation and training protocol).

### Experimental Phase

During the experiments, feeders were obscured with non-perforated paper. Each bird was tested in four different experimental set-ups, a choice between: 1) a provisioned and an empty feeder marked with a combination of corresponding visual and olfactory cues (banana or almond); 2) a provisioned and an empty feeder marked with only olfactory cues (banana or almond); 3) a provisioned and an empty feeder marked only with visual cues; and 4) a provisioned and an empty feeder without any cues (Figure 1B). To avoid side bias, we repeated each trial with feeder positions reversed, and tested all birds in all feeder combinations. Thus, each bird participated in twelve trials. Birds participated in a single trial per day, to ensure choices were independent and unbiased by recent experience, and to avert trial fatigue. All trials were conducted between 9am-3pm.

**Figure 1.**
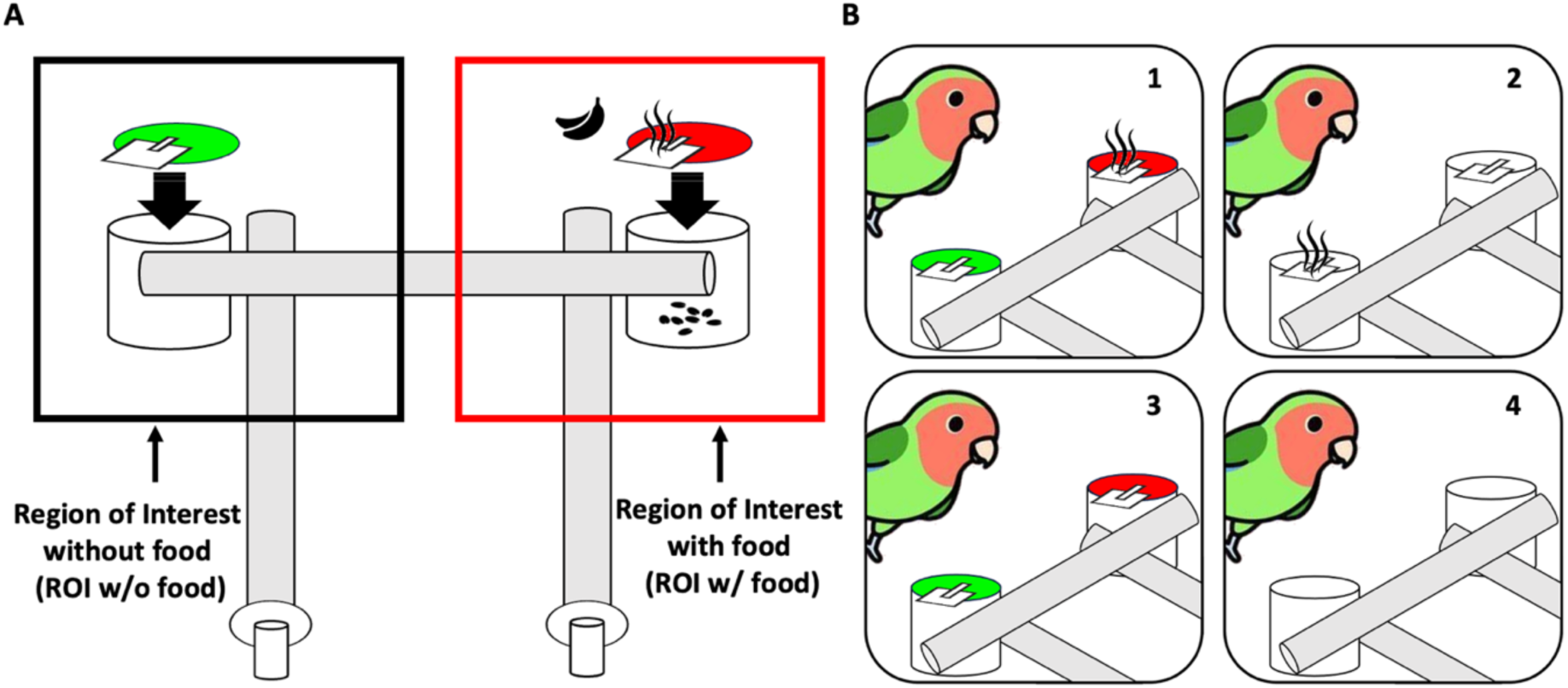
A) A schematic representation of the experimental design, including both the visual cue (red/green paper) and/or olfactory cue (either banana scent on a provisioned feeder or almond scent on an empty feeder). The region of interest was defined for each feeder; B) Each bird participates in four treatments, 1: both visual and olfactory cues; 2: olfactory cue only; 3: only visual cue; 4: no cues.

### Behavioural analysis

We recorded bird behaviour using a video camera (STARCAM CB71 Mini Battery IP Camera) placed above the cage (Figure S1). We defined two 12cm × 12cm regions of interest (ROI) centred around either feeder (ROI_food_: ROI around a feeder_food_; ROI_w/o food_: ROI around a feeder_w/o food_) (Figure 1A). We classified behaviours into ‘investigation’ (i.e., head turns, body turns, preening, and wing stretching) and ‘tearing’ (i.e., bird using its beak to tear a hole in the paper cover). To standardise video length between individuals and trials, we trimmed videos to start once the bird entered the field of vision, and to end once the bird chose one of the feeders (i.e., tearing a hole in the paper and putting its head into the feeder). If birds failed to do so within 1 hour, no choice preference was recorded for that trial and the video was not included in downstream analysis. Birds that tore away the paper covers of both feeders, but without making a clear choice (i.e., putting their head into neither feeder), were assigned a choice based on the duration spent in each feeder ROI, time spent investigating each feeder, and time spent tearing at each feeder prior. Videos were recorded at a resolution of 1920 × 1080 pixels, but were down-sampled to 1280 × 720 pixels, with a bit-rate of 1000 bps at 30 frames per second (fps,) to facilitate further computational analysis (Mathis & Warren, 2018).

### Machine learning of pose estimation and behavioural classification

We used DeepLabCut (2.2.0.3; Mathis et al., 2018) for markerless tracking of the relative positions of seven body parts (left eye, right eye, crown, beak, left nape, right nape, and back centre; Figure S2) for pose-estimation. Simple Behavioural Analysis (SimBA) (0.89.9; Nilsson et al., 2020) was used for supervised machine learning of behavioural predictive classification to quantify recorded behaviours automatically. A random subset of 3000 labelled frames from 50 videos, taken during different experiments featuring different individuals, was used for network training.

We trained a ResNet-50-based neural network (He et al. 2016; a convolutional neural network, up to 50 layers deep), set with default parameters, and using the maximum of 10,300,000 training iterations. Validation, based on a single shuffle (to normalise data) gave a test error of 3.15 pixels and a train error of 2.66 pixels (with p-cut-off =0.95). We used a p-cut-off of 0.95 to condition the X, Y coordinates for future analysis. Network training was performed in the Google Colab Pro environment (Carneiro et al., 2018) with NVIDIA Tesla T4/P100 GPUs. The trained network was applied to analyse all videos, yielding pose tracking files for subsequent analysis. Figure 2A shows the pose-estimation analytical procedure. The video and the tracking file of each bird were input into SimBA to produce behavioural classifiers (Nilsson et al., 2020).

**Figure 2.**
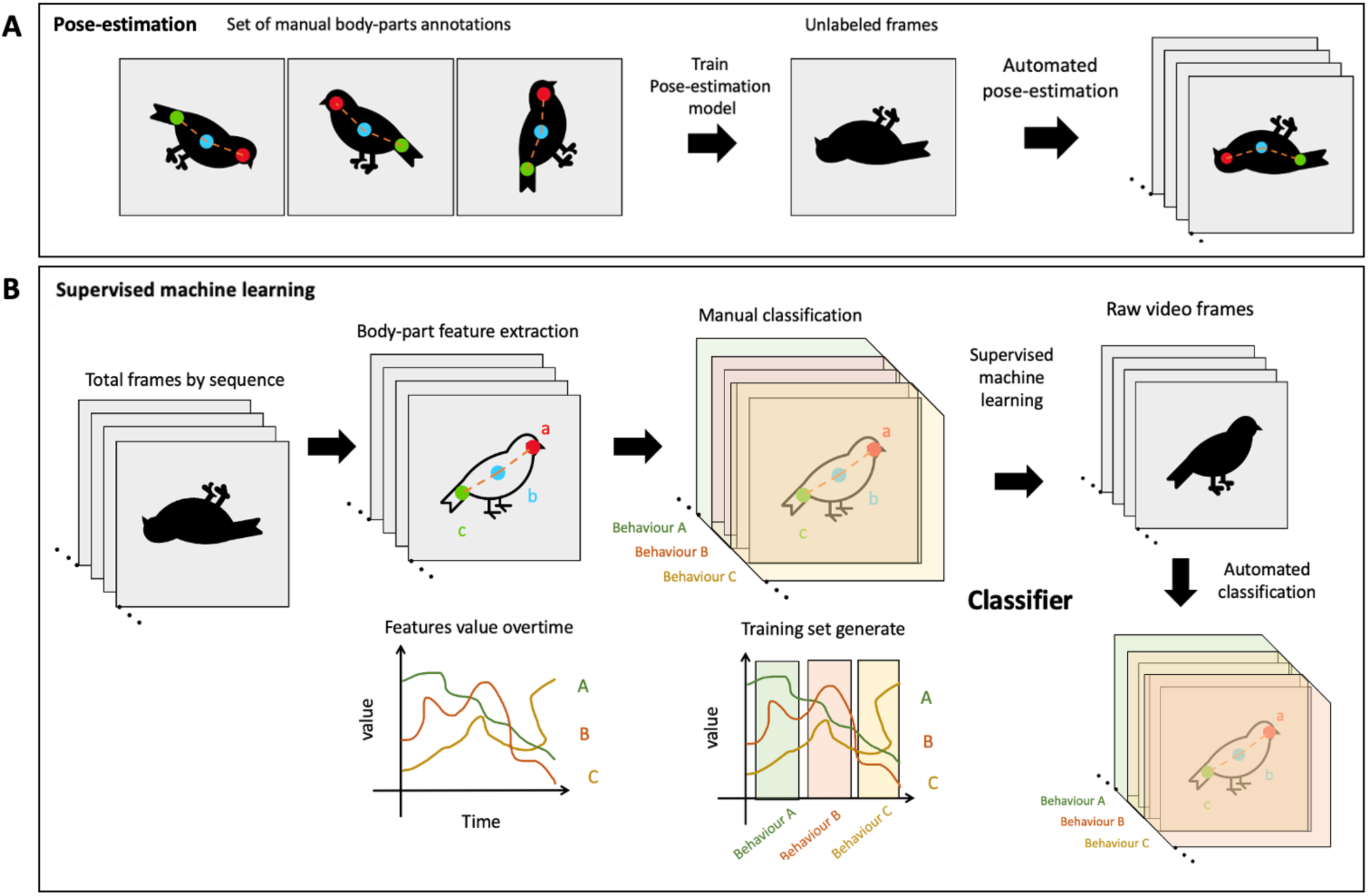
A) Pose-estimation algorithms used to track animal body parts based on manual annotations in a set of training videos. The model marked corresponding body parts based on input data, used to analyse raw videos. B) Supervised machine learning trained the classifier with manually defined behaviours. The trained classifier then detected these key data in new video and identified these behaviours according to this learning algorithm data. Figure modified from von Ziegler et al. (2021).

Next, we took pose-estimation data, extracted from the DeepLabCut procedure, standardised for relative body-position distance (pixels/mm) movement, angles, areas, and path metrics and their deviations and rank for individual frames and across rolling windows, along with time (fps) standardisation, to which we applied the SimBA in-built event logger (using FFmpeg to display individual frames alongside extracted video) to annotate the presence/absence of each behaviour within each ROI, the total duration of each behaviour, and the time when the bird entered/exited either ROI. Behaviour classifier models separated data and eigenvalues into different classes (e.g., absence or presence) by applying intuitive random forest classifier algorithms (trained using default hyperparameters; Figure 2B) to these raw video data to obtain the behavioural dataset.

Each classification model was evaluated based on calculating its precision, recall and F1 curve scores after 5-fold shuffle cross-validations on 20% of the datasets annotated in the SimBA event logger, and by testing each classifier on the un-shuffled, correctly annotated behavioural event annotations in the training data set. We generated classifier learning curves that indicated how the inclusion of further logged behavioural events affected classifier performance. We also evaluated F1-scores for learning curves after performing 5-fold cross-validations using 1, 25, 75 and 100% of the shuffled data sets to predict the classified behaviours on 20% of the datasets. Precision-recall curves were generated to visualise how classifiers can be titrated to balance classification sensitivity against specificity across different discrimination thresholds, which we used to set the optimal discrimination threshold for ‘investigation’ at 0.52 and ‘tearing’ at 0.5625 (see Figure S3 for illustration of the workflow for machine learning, pose estimation and behavioural classification).

Mean precision, recall, and F1-scores for the presence/absence of behaviours following the 5-fold shuffle cross-validation of each classifier are presented in Figure S4. Classification performance for the presence of behaviours as measured by F1 were 0.736 and 0.781, precision was 0.713 and 0.735, and recall was 0.758-0.834; the classification performance scores for the absence of behaviours were slightly higher (Figure S4C). Five-fold cross-validation learning curves using 1-100% of these annotated data (Figure S4A) showed that the number of annotated images correlated positively with the F1-score. Precision-recall curves (Figure S4B) indicated optimal classifier performance for different classifications, as measured by F1-score at discrimination thresholds between 0.52 and 0.56. We used SimBA to correct gross pose-estimation tracking inaccuracies based on impossible locations and movements of body parts.

From the pose-estimation model and the behavioural classifier, we calculated the proportion of each response variable (i.e., time spent on ‘investigation’ and ‘tearing’, number of times the bird entered either ROI, and total time spent in each ROI; but excluding choice result and choice latency, see Table 1) in each trial by dividing the total time spent on each behaviour by the total video duration time. Choice latency was first calculated as the total video duration minus the total time the bird spent in each ROI, and standardized as a proportion from 0 to 1 for better visualisation. No data were standardised before conducting any statistical analyses.

**Table 1.** Definition of response variables.

| **Data collected** | **Type of data** | **Definition** | **Collection method** |
| --- | --- | --- | --- |
| Choice result | Bivariate | The feeder first opened entirely, into which the bird sticks its head | Recorded by observer |
| No. of ROI entry | Count | Each event = the entry of the crown of the bird's head into the ROI | Automatically calculated using SimBA |
| Investigation time | Continuous | The duration of time the bird spends investigating the feeder within each ROI, including body turns, head turns, preening, and wing stretching, but without touching the feeder directly. | Automatically calculated using SimBA |
| Choice latency | Continuous | Calculated as total video duration minus the total time the bird spent in each ROI; an indicator of choice decision hesitancy reflecting time to switch between ROIs | Automatically calculated using SimBA |
| Tearing time | Continuous | The duration of time the bird performing tearing behaviour on the feeder within each ROI, recorded from when the bird touches the feeder directly | Automatically calculated using SimBA |
| Total time spent | Continuous | Total time spent in each ROI during the entire video | Automatically calculated using SimBA |

Choice latency (i.e., when birds were not in either ROI) was analysed equally for ROI_food_ and ROI_w/o food_. In contrast, the frequency with which birds entered either ROI, and the total time spent in these ROIs, were analysed based on the ROI_food_. Consequently, latency was reciprocally proportionate between the two ROIs, and analysed as a total for each trial, whereas investigation time and tearing were analysed separately for each ROI, as these data were not reciprocally proportionate.

### Statistical analyses

Statistical analyses were performed using R (version 4.2.0) with RStudio. We used the "GLMMTMB" package for generalised linear mixed models (Magnusson et al., 2017). Two identical models were performed, one for each olfactory cue, the ‘banana model’ and the ‘almond model’. In both models, N designates the no-cues treatment group ; O that only olfactory cues were presented; V that only visual cues were presented, and B that both sensory cues were presented. We also analysed the complete suite of behaviours recorded during each trial to test for a combinatory effect between visual and olfactory cues.

Visual cues (yes/no) and olfactory cues (yes/no for either banana or almond) were included as fixed effects with an interaction term. Individual identity and trial number were included as random effects, with the side of the feeder_food_ included as a random slope. The significance of successfully selecting the feeder_food_ (‘success rate’) was determined using binomial statistics. Multiple pairwise comparisons were subjected to Benjamini-Hochberg correction. Critical alpha values were set at *p*<0.05 for all analyses, unless stated otherwise.

We used the “DHARMa” and “performance” packages to check if model assumptions were met (Lüdecke et al., 2021). We checked normality among model residuals and each random effect with Shapiro-Wilk tests and qqplot, and checked for homogeneity of variances, linearity of fitted value and residuals, and collinearity of the variance inflation factor (VIC) using Levene’s tests and Bartlett’s tests. We used boxplots to identify influential outliers. If the model residual did not fit normality, a Box-Cox transformation was applied to these data (Box & Cox, 1964). Although results were similar if outliers were included, outliers were excluded if they significantly affected model homogeneity of the variance to ensure model assumptions were met.

## Results

### Behavioural responses: selecting the feeder that contained the food reward

We analysed a total of 296 videos. Treatment cues affected the success rate at which birds correctly chose the feeder_food_ (Figure 3): Overall, if only visual cues were presented (Figure 1 B3), success rate was 96.15% (50/52), significantly better than expected by chance (*p*<0.001); if no cues were presented (Figure 1 B4), overall success rate was 47.92% (23/48), i.e., not significantly different from random choice (*p*=0.11). If a positive visual cue indicating food (i.e., red paper) was paired with banana scent (Figure 1 B1), success rate was 87.23% (41/47, p<0.001); if a negative visual cue (green paper) was paired with almond scent, success rate was 98% (49/50, *p*<0.001). If, however, only scent cues were presented (Figure 1 B2), success rate dropped to 36.54% (19/52, *p*=0.017) for banana scent and 61.7% (29/47, *p*=0.03) for almond scent (Figure 3).

**Figure 3.**
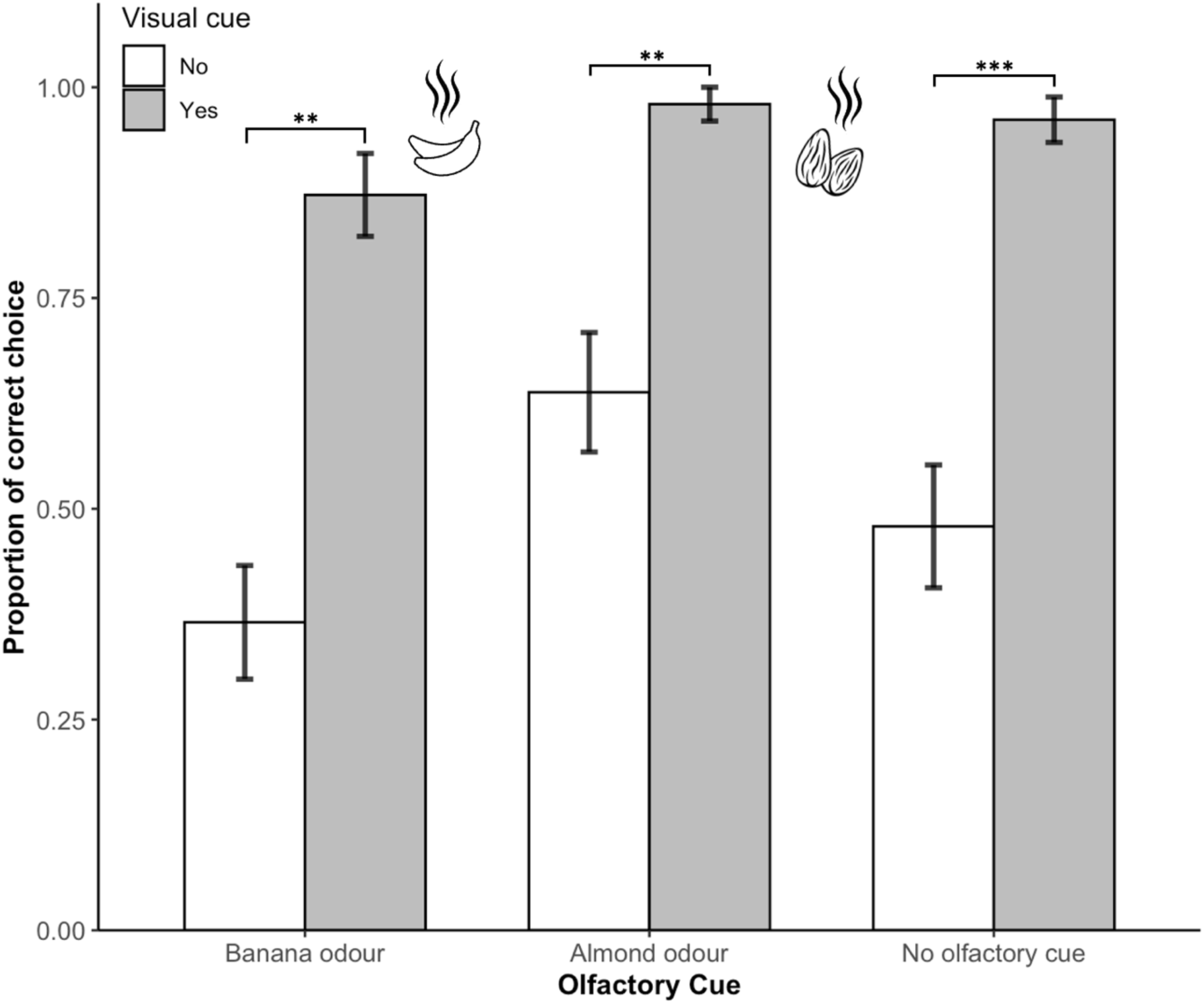
Choices made based on olfactory and visual cues. The y-axis represents the proportion of times the bird chose the feeder containing food in each treatment; the x-axis and legends indicate each treatment group. Error bars gives the standard error of the mean. Significant post-hoc comparisons are denoted by asterisks (*p<0.05; **p<0.01; ***p<0.001).

#### 1) Feeder choice

Only visual cues affected feeder choice significantly (banana model ANOVA: *χ^2^_1,196_*=18.16, *p*<0.001; almond model ANOVA: *χ^2^_1,194_*=27.97, *p*<0.001), with no significant interactions in either model (Table. 2 & 3; Figure 3). Post-hoc model comparison showed that feeder choice differed only when visual cues were present, i.e., between treatment groups N and V (banana model: t_196_= -3.19, *p*<0.01; almond model: t_194_= -4.25, *p*<0.001); N and B (banana model: t_196_= -2.36, *p*<0.05; almond model: t_194_= -3.79, *p*<0.001); V and O (banana model: t_196_= 3.66, *p*<0.01; almond model: t_194_= 3.38, *p*<0.01); and B and O (banana model: t_196_= -2.92, *p*<0.01; almond model: t_194_= -3.15, *p*<0.01).

#### 2) Number of ROI_Food_ entries

The number of ROI_Food_ entries prior to the bird choosing a feeder was also only affected by visual cues (banana model: ANOVA: *χ^2^_1,195_*=64.94, *p*<0.001; almond model: ANOVA: *χ^2^_1,195_*=12.92, *p*<0.001) with no significant interactions in either model (Table. 2 & 3; Figure 4). Post-hoc comparison showed that ROI_Food_ entries differed significantly between treatment groups with and without visual cues (banana model: N and V: t_195_= -5.33, *p*<0.001; N and B: t_195_= -5.14, *p*<0.001; V and O: t_195_= 6.30, *p*<0.001; B and O: t_195_= -6.05, *p*<0.001; almond model: N and V: t_193_= -2.90, *p*<0.05; N and B: t_193_= - 2.91, *p*<0.05; V and O: t_193_= 2.17, *p*<0.05; B and O: t_193_= -2.18, *p*<0.05).

**Figure 4.**
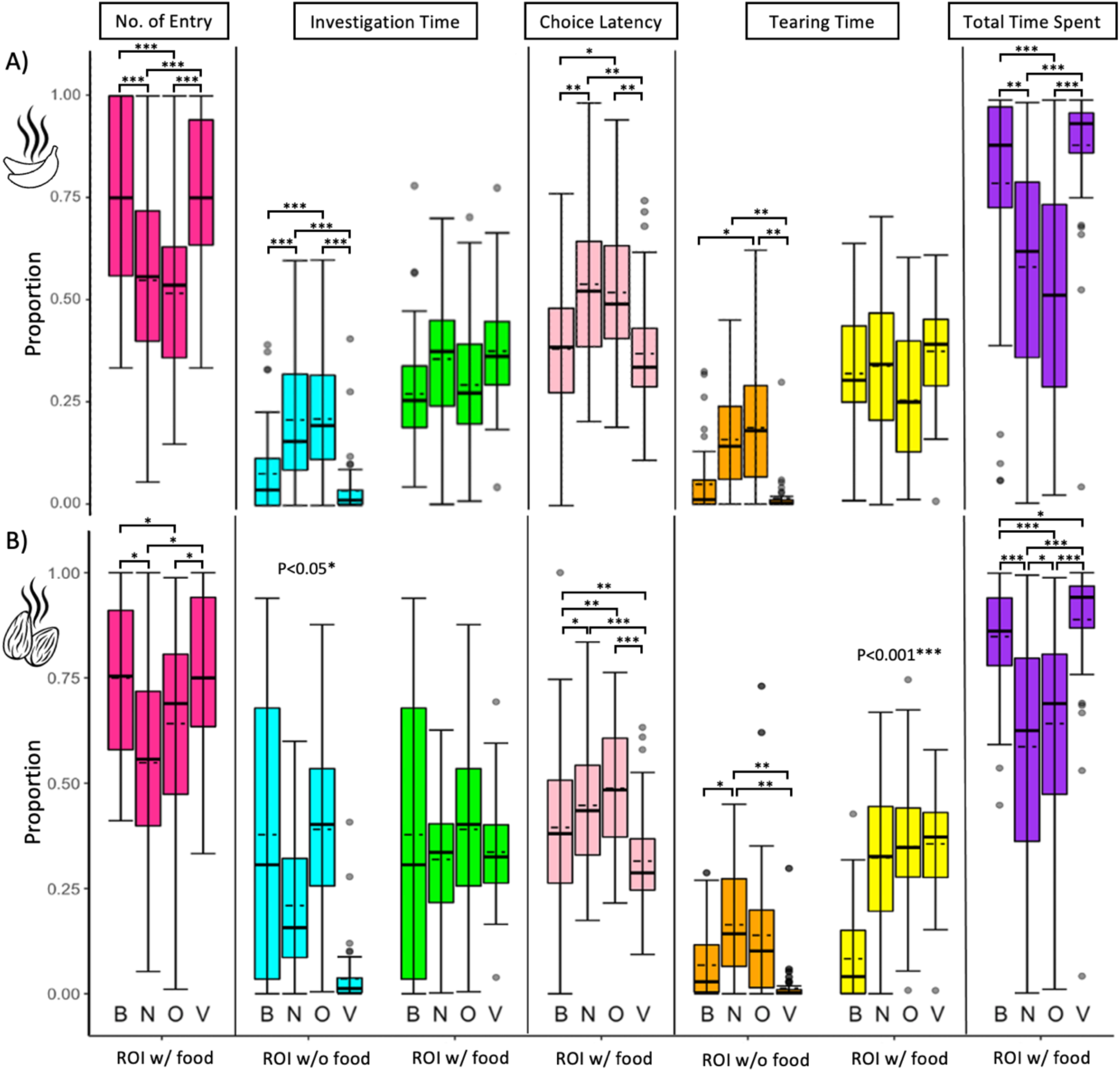
Box and whisker plots showing variation in response variables among treatments and their relation to each ROI. A) Banana scent as the olfactory cue. B) Almond scent as the olfactory cue. Each response variable is indicated at the top of the figure, and each box reflects the result of a different treatment as indicated by the capital letter at the bottom of each panel (B: Both cues; N: No cues; V: visual cue only; O: olfactory cue only). The horizontal bar in each box represents the median. The dashed line indicates the mean. Dots represent outliers. ROIs (with or without food) were indicated at the bottom of the figure. Significant p-values are denoted by asterisks (*p<0.05; **p<0.01; ***p<0.001) based on post-hoc results. Significant p-values without parentheses indicate that an interaction occurred between treatment groups.

#### 3) Investigation time

Time spent investigating either ROI_Food_ or ROI_w/o Food_ was affected by experimental treatments. For the banana model, only visual cues significantly affected the time spent investigating the ROI_w/o Food_ (ANOVA: *χ^2^_1,175_*=75.33, *p*<0.001), with no significant interactions between visual and olfactory cues (Table 2; Figure 4A). Post-hoc comparison showed significant differences between treatment groups N and V (t_175_=7.29, *p*<0.001); N and B (t_175_=4.72, *p*<0.001); V and O (t_175_= -7.62, *p*<0.001); and B and O (t_175_=4.98, *p*<0.001). The time spent investigating in the ROI_Food_ was only affected significantly by olfactory cues (ANOVA: *χ^2^_1,195_*=5.93, *p*<0.05) without significant interactions between visual and olfactory cues (Table 2; Figure 4A). Post-hoc comparison found no significant differences between treatment groups.

**Table 2.**
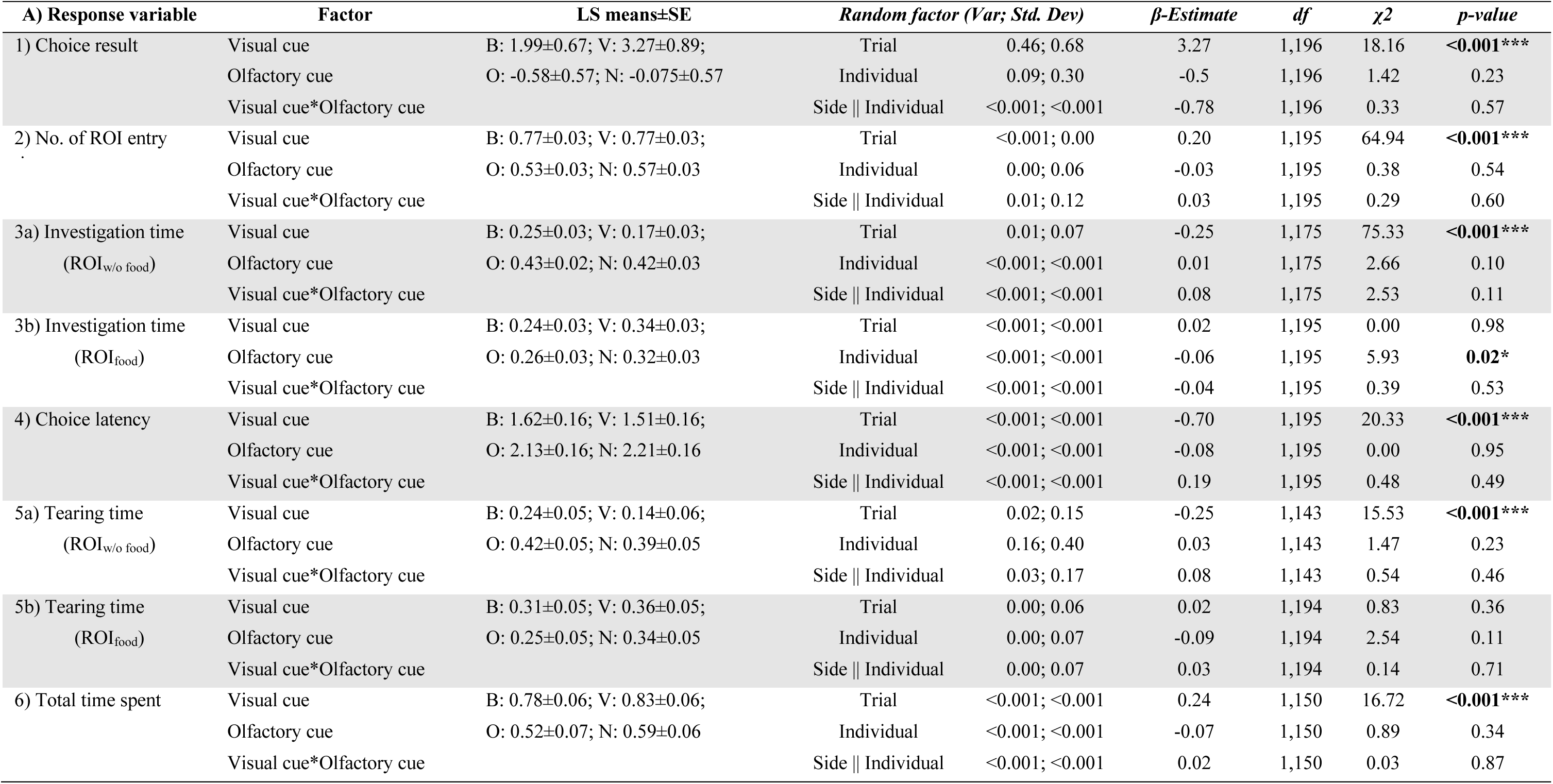
Bird responses to four different treatments (using banana flavour): B: both cues; V: only visual cue; O: only olfactory cue; N: no cues presented. Significant p-values are in bold and denoted by asterisks (*p<0.05; **p<0.01;***p<0.001), post-hoc results also shown in bold if significant. ROI_food_: Region of interest with food. ROI_w/o food_: Region of interest without food.

For the almond model, visual cues significantly affected time spent investigating the ROI_w/o Food_ (ANOVA: *χ^2^_1,165_*=40.51, *p*<0.001). The interaction between visual and olfactory cues significantly affected time spent investigating the ROI_w/o Food_ (ANOVA: *χ^2^_1,165_*= 11.62, *p*<0.05) (Table 3; Figure 4B), but there were neither significant effects nor a significant interaction between visual and olfactory cues (Table 3; Figure 4B); therefore, post-hoc comparison found no significant differences between treatment groups.

**Table 3.** Bird responses to four different treatments (using almond flavour): B: both cues; V: only visual cue; O: only olfactory cue; N: no cues presented. Significant p-values are in bold and denoted by asterisks (*p<0.05; **p<0.01;***p<0.001), post-hoc result also shows in bold if significant. ROI_food_: Region of interest with food. ROI_w/o food_: Region of interest without food.

| B) Response variable | Factor | LS means±SE | Random factor (Var; Std. Dev) |  | β-Estimate | df | χ2 | p-value |
| --- | --- | --- | --- | --- | --- | --- | --- | --- |
| 1) Choice result | Visual cue | B: 3.90±1.02; V: 3.23±0.73; | Trial | <0.001; <0.001 | 3.32 | 1,194 | 27.97 | <0.001*** |
|  | Olfactory cue | O: 0.58±0.32; N: -0.08±0.30 | Individual | 0.20; 0.45 | 0.66 | 1,194 | 2.76 | 0.1 |
|  | Visual cue*Olfactory cue |  | Side \|\| Individual | <0.001; <0.001 | 0.01 | 1,194 | 0.00 | 0.99 |
| 2) No. of ROI entry times | Visual cue | B: 0.97±0.01; V: 0.97±0.01; | Trial | <0.001; <0.001 | 0.04 | 1,193 | 12.92 | <0.001*** |
|  | Olfactory cue | O: 0.94±0.01; N: 0.93±0.01 | Individual | <0.001; <0.001 | 0.01 | 1,193 | 0.27 | 0.60 |
|  | Visual cue*Olfactory cue |  | Side \|\| Individual | <0.001; <0.001 | -0.01 | 1,193 | 0.25 | 0.62 |
| 3a) Investigation time<br>(ROI<sub>w/o food</sub>) | Visual cue | B: 0.27±0.04; V: 0.14±0.03; | Trial | <0.001; 0.02 | -0.26 | 1,165 | 40.51 | <0.001*** |
|  | Olfactory cue | O: 0.33±0.03; N: 0.40±0.03 | Individual | <0.001; <0.001 | -0.07 | 1,165 | 0.01 | 0.94 |
|  | Visual cue*Olfactory cue |  | Side \|\| Individual | <0.001; <0.001 | 0.21 | 1,165 | 11.62 | 0.037* |
| 3b) Investigation time<br>(ROI<sub>food</sub>) | Visual cue | B: 0.39±0.05; V: 0.34±0.05; | Trial | <0.001; <0.001 | -0.02 | 1,140 | 0.08 | 0.78 |
|  | Olfactory cue | O: 0.45±0.06; N: 0.36±0.05 | Individual | <0.001; <0.001 | 0.04 | 1,140 | 2.02 | 0.15 |
|  | Visual cue*Olfactory cue |  | Side \|\| Individual | <0.001; <0.001 | 0.08 | 1,140 | 0.60 | 0.44 |
| 4) Choice latency | Visual cue | B: 1.89±0.16; V: 1.51±0.16; | Trial | 0.02; 0.15 | 1.51 | 1,191 | 20.46 | <0.001*** |
|  | Olfactory cue | O: 2.36±0.16; N: 2.21±0.16 | Individual | 0.19; 0.44 | 0.15 | 1,191 | 4.15 | 0.04* |
|  | Visual cue*Olfactory cue |  | Side \|\| Individual | 0.07; 0.26 | 0.23 | 1,191 | 0.78 | 0.38 |
| 5a) Tearing time<br>(ROI<sub>w/o food</sub>) | Visual cue | B: 0.25±0.04; V: 0.15±0.04; | Trial | 0.00; 0.05 | -0.24 | 1,131 | 16.27 | <0.001*** |
|  | Olfactory cue | O: 0.37±0.04; N: 0.39±0.04 | Individual | <0.001; 0.02 | -0.02 | 1,131 | 0.62 | 0.43 |
|  | Visual cue*Olfactory cue |  | Side \|\| Individual | <0.001; 0.03 | 0.12 | 1,131 | 1.95 | 0.16 |
| 5b) Tearing time<br>(ROI<sub>food</sub>) | Visual cue | B: 0.08±0.03; V: 0.35±0.03; | Trial | <0.001; <0.001 | 0.04 | 1,182 | 14.22 | <0.001*** |
|  | Olfactory cue | O: 0.34±0.03; N: 0.31±0.03 | Individual | <0.001; <0.001 | 0.02 | 1,182 | 19.88 | <0.001*** |
|  | Visual cue*Olfactory cue |  | Side \|\| Individual | <0.001; <0.001 | -0.29 | 1,182 | 26.57 | <0.001*** |
| 6) Total time spent | Visual cue | B: 0.73±0.04; V: 0.83±0.04; | Trial | <0.001; 0.03 | 0.26 | 1,172 | 62.26 | <0.001*** |
|  | Olfactory cue | O: 0.46±0.04; N: 0.57±0.04 | Individual | 0.01; 0.08 | -0.11 | 1,172 | 9.16 | <0.01** |
|  | Visual cue*Olfactory cue |  | Side \|\| Individual | 0.00; 0.05 | 0.02 | 1,172 | 0.11 | 0.74 |

#### 4) Choice latency

For the banana model, only visual cues significantly affected choice latency (ANOVA: *χ^2^_1,195_*=20.33, *p*<0.001), with no significant interactions between visual and olfactory cues (Table 2; Figure 4A). Post-hoc comparison found a significant difference between treatment groups N and V (t_195_= 3.68, *p*<0.01); N and B (t_195_= 3.11, *p*<0.01); V and O (t_195_= -3.27, *p*<0.01); and B and O (t_195_= 2.69, *p*<0.05).

For the almond model, however, both visual and olfactory cues significantly affected choice latency (ANOVA: visual *χ^2^_1,191_*=20.46, *p*<0.001; olfactory: *χ^2^_1,191_*=4.15, *p*<0.05), with no significant interaction between visual and olfactory cues (Table 3; Figure 4B). Post-hoc comparison found a significant difference between treatment groups N and V (t_191_= 3.84, *p*<0.001); V and O (t_191_= -4.64, *p*<0.001); and B and O (t_191_= 2.55, *p*<0.05).

#### 5) Tearing time

Both models evidenced different tearing times for ROI_Food_ and ROI_w/o Food_. For the banana model, only visual cues significantly affected tearing time in the ROI_no Food_ (ANOVA: *χ^2^_1,143_*=15.53, *p*<0.001), without significant interaction between visual and olfactory cues (Table 2; Figure 4A). Post-hoc comparison found a significant difference between treatment groups N and V (t_195_=3.29, *p*<0.01); V and O (t_195_= -3.67, *p*<0.01); and B and O (t_195_=2.29, *p*<0.05). The time spent tearing in the ROI_Food_ was not affected by cues, with no significant interactions between visual and olfactory cues (Table 2; Figure 4A). Post-hoc comparison found no significant differences between treatment groups.

For the almond model, only visual cues significantly affected tearing time in the ROI_w/o Food_ (ANOVA: *χ^2^_1,131_*=16.27, *p*<0.001), without significant interaction between visual and olfactory cues (Table 3; Figure 4B). Post-hoc comparison found a significant difference between treatment groups N and V (t_131_=3.82, *p*<0.01); B and N (t_131_=2.27, *p*<0.05); and V and O (t_131_= -3.45 *p*<0.01). For the ROI_Food_, there was a significant interaction between the visual and olfactory cues in tearing time (Table 3; Figure 4B).

#### 6) Total time spent

The total time spent in ROI_Food_ was affected by visual cues (banana model: ANOVA: *χ^2^_1,150_*=16.72, *p*<0.001; almond model: ANOVA: *χ^2^_1,172_*=62.26, *p*<0.001), and olfactory cues were also significant in the almond model (ANOVA: *χ^2^_1,172_*=9.16, *p*<0.01) (Table. 2 & 3; Figure 4). Post-hoc comparison found a significant difference between treatment groups with and without visual cues, i.e., between treatment groups N and V (banana model: t_146_= -4.15, *p*<0.001; almond model: t_172_= -5.13, *p*<0.001); N and B (banana model: t_146_= -3.22, *p*<0.01; almond model: t_172_= -3.27, *p*<0.001); V and O (banana model: t_146_= 5.16, *p*<0.001; almond model: t_172_= 8.03, *p*<0.001); and B and O (banana model: t_146_= - 4.21, *p*<0.001; almond model: t_172_= -6.00, *p*<0.001). For the almond model, total time spent was also different between the treatment groups N and O (t_172_= -2.27, *p*<0.05) and B and V (t_172_= 2.02, *p*<0.05).

## Discussion

Lovebird feeder choice was guided almost exclusively by visual cues. If presented with a visual cue, they were significantly more likely to choose the feeder_food_, to spend more time in the ROI_Food_, and to spend less time hesitating (latency) on which feeder to choose. They also performed significantly less ‘investigation’ and ‘tearing’ behaviours at the feeder_w/o food_ when a visual cue was presented. In contrast, neither banana (positive cue) nor almond (negative cue) odour influenced their feeder choices (Figure 3). In fact, their foraging behaviour when only olfactory and no visual cues were presented was very similar to when no cues were presented. They also spent a similar duration investigating and tearing at a visually identifiable feeder_food_, irrespective of whether olfactory cues were presented.

From this we deduce that lovebirds did not utilise the olfactory cues we provided to locate their food, but instead their decision making and processing time were informed solely by visual cues. We also found that, once lovebirds established that a feeder contained no food, they gave it little subsequent attention. This is consistent with optimal foraging theory, which posits that species should adopt the most economically advantageous foraging patterns (Pyke et al., 1977).

Various previous studies have established the importance of visual foraging cues for birds, and so our results are not unexpected. Most fruits, berries, and seeds have evolved distinctive colours with wavelengths easily distinguished by birds (Lei et al. 2021), and strong visual ability enables insectivorous birds to detect prey directly (Zvereva et al. 2019), and to detect even subtle differences in the reflective spectrum produced by herbivore-damaged trees (Mäntylä et al. 2020). The importance of olfaction in avian sensory perception has also been demonstrated across species and behaviours. For instance, plants suffering herbivory engage in a tritrophic response, releasing volatile secondary allelochemical metabolites to attract predators to remove these grazers (Mäntylä et al., 2004). Nevitt et al. (1995) found that Procellariiform seabirds are attracted to the dimethyl sulphide scent produced by phytoplankton in response to zooplankton grazing, resulting in the seabirds consuming these herbivores. Rubene et al. (2019), however, showed experimentally that although herbivore-induced plant volatiles attracted various insectivorous bird species, these olfactory cues were still secondary to visual cues in avian foraging decisions. Importantly, and contrary to other studies (e.g., Mäntylä et al., 2020; Potier et al., 2019; Rubene et al., 2019), in lovebirds we did not find any evidence that olfaction was acting synergistically with vision. This suggests that while vision is augmented by scent in some bird species, this synergism has not acted as a selection pressure during the evolution of lovebird sensory perception (Steiger et al., 2010).

Unlike most other parrot species, known to have a keen sense of smell (e.g., yellow-backed chattering lories *Lorius garrulus flavopalliatus*: Roper 2003; kakapo *Strigops habroptilus*: Hagelin 2004; kea and kaka: Gsell, 2012, budgerigars: Zhang et al., 2010), rosy-faced lovebirds live in arid habitats, with high temperatures, intense sunlight, and low humidity; conditions not conducive to the persistence of odour molecules that evaporate/denature quickly (Ndithia & Perrin, 2009b). In this habitat, however, edible plants typically have high visual salience. Furthermore, unlike other parrot species that may use their keen sense of smell to locate ripe fruit (e.g., the nocturnal kakapo [*Strigops habroptila*]: Gsell et al., 2012; macaws: Hernández et al., 2022), lovebirds rarely feed on fruits in the wild (Ndithia & Perrin, 2009a). Olfactory ability is reflected in the olfactory bulb to brain ratio, which, along with OR gene numbers, is likely affected by ecological factors and life-history adaptations (e.g., nocturnal lifestyle: Gsell et al., 2012; trophic niche: Toda et al. 2021). In birds, olfactory bulb to brain ratio correlates positively with the estimated total number of OR genes (Steiger et al. 2010; Khan et al. 2015), although parrots generally have relatively small ratios (Corfield et al. 2015). Since most *Agapornis* species, including *A. roseicollis* and their common ancestor, are granivorous (Huynh et al. 2023), their olfactory ability (or use of olfaction in foraging) is likely more limited compared to parrot species that have evolved strong olfactory ability. Therefore, that lovebirds do not associate banana scent with foraging success may be due to a lack of association, a lack of olfactory receptors expressed in sensory neurons within the olfactory epithelium, or due to the loss of functional olfactory receptor genes able to detect certain odour cues (i.e., almond) (Steiger et al., 2010).

Nevertheless, although banana or almond odour cues did not improve rosy-faced lovebird foraging decisions, this does not necessarily infer insensitivity to all odours. Other studies have found that foraging birds can be much more sensitive to one odour than another (Kelly & Marples, 2004). For instance, birds can be highly sensitive to irritant odours (e.g., 2-methoxy-3-sec butyl pyrazine; 2-methoxy-3-isobutyl pyrazine; mint) that stimulate not only the olfactory nerve but also trigger the trigeminal nerve, perceived as pain (Müller-Schwarze, 2006). Further studies will need to ascertain if this is true also for *A. roseicollis*.

Furthermore, while our investigation suggests that rosy-faced lovebirds do not rely on olfaction while foraging, olfaction could still be important in non-foraging contexts. For example, birds use scent and olfaction in manifest social contexts (Liu, 2022), such as determining sex through the scent of uropygial gland secretion (Amo et al., 2012), chemo-signalling reproductive status (Caro et al., 2015), and even using olfaction to detect MHC compatibility (Leclaire et al., 2017). Humboldt penguins (*Spheniscus humboldti*) use odour to recognize familiar and related conspecifics (Coffin et al. 2011); female zebra finches (*Taeniopygia guttata*) and Bengalese finches (*Lonchura striata var. domestica*) use odour cues to recognize their own nests (Krause & Caspers, 2012); house finches (*Carpodacus mexicanus*) can detect olfactory cues from predatory and non-predatory mammalian faeces (Roth II et al., 2008). As rosy-faced lovebirds are highly social, future research should test whether they use olfaction in non-foraging behavioural contexts.

Generally, the ability of a species to adapt to captivity and new objects directly affects the success of behavioural experiments (Lambrechts et al., 1999), where neophobia can constrain research on bird perception and cognition (Greenberg, 2003). Lovebirds learnt the layout and principles of our experimental set-up rapidly during the habituation phase, forming a positive association between colour and food. This implies that they are intelligent enough to participate in this type of empirical investigation, comparable to other parrot species (Auersperg and von Bayern, 2019). Future work should investigate the intelligence of lovebirds, particularly their ability to recognize objects, as established for larger parrots.

From a technical perspective, the advanced machine-learning analysis we applied substantially overcame the limitations of having a human observer annotate, identify, record, and interpret relevant behavioural changes in real time, where it requires 22 person-hours to annotate a 1-hour video with frame-by-frame precision (Jhuang et al., 2010). Furthermore, having multiple people annotate recordings can result in inter-observer error (von Ziegler et al., 2021). Machine-learning saved us approximately 503 person-hours of video annotation (with associated staff costs). Importantly, our approach did not require any motion capture (Microelectromechanical systems, MEMS) markers (Mishra & Kiourti, 2021) to be attached to the birds (Won et al., 2021), which can cause discomfort and distress, compromising psychological, behavioural, and physiological data quality (Sneddon, 2017). Markerless motion capture technology (Nakano et al., 2020), combined with deep learning computational approaches, such as the convolutional neural networks (CNN) applied by the DeepLabCut open-source software, (Mathis et al., 2018), thus has significant potential to advance recording and processing of animal pose-estimation in behavioural studies (Labuguen et al., 2021; Li et al., 2023).

In conclusion, ours is the first study to apply deep-learning techniques to expand on the role of avian olfaction in optimal foraging, involving a non-standard avian model species, providing a standardised protocol for future behaviour studies. That lovebirds did not use olfaction to detect food, relying instead on visual perception, highlights how sensory processing sensitivity is strongly related to the environmental conditions in which each species lives, affecting their sensory neurophysiology (Corfield et al., 2015) and genetics (Khan et al., 2015; olfactory receptor gene repertoires—Steiger et al., 2010), which ultimately shapes their life-history evolution (Driver & Balakrishnan, 2021; Steiger et al., 2010) and personality traits (Dingemanse & Réale, 2015).

## Ethics

All procedures were approved by the University of Hong Kong Committee on the Use of Live Animals in Teaching and Research (CULATR; approval number: 5883-21), and under a Department of Health Animal (Control of Experiments) Ordinance Chapter 340 permit ((21-1146) in DH/HT&A/8/2/3 Pt.32).

## Acknowledgements

We thank Yiu Siu, Ellen Sai Nam Lo, Tat Sing Ngai, and Mei Ying Wu for animal care and husbandry.

## Supplementary Materials

## Supplementary methods

### Habituation and Training

Prior to our experimental testing, we conducted a habituation phase, during which birds were allowed to forage and familiarize themselves with the experimental set-up. To train the birds to seek the non-visible food rewards obscured by the paper cover, we first perforated the paper over each feeder bowl with a small hole so that the food was partially visible. Birds thus learned to remove the paper to access the food. We ascertained that all study subjects had understood the principal by monitoring that the birds i) looked through the hole at the food first before they removed the paper cover, ii) removed the paper cover on provisioned feeders, and iii) lost interest in the feeder once they had emptied it of food. Success rate in this pilot experiment (n=37) was 0.81 (binomial test; *p*<0.001), indicating that bird behaviour in our experiments was motivated by food, which ensured the reliability of our observations. During training, feeder positions were assigned randomly to each side of the perch apparatus to avoid side bias. During the subsequent associative learning phase, we continued using perforated papers and the birds were trained to associate the visual red/green coloured paper cues or olfactory banana/almond odour cues with the presence/absence of a food reward, for approximately 10 rounds for either cue.

## Supplementary figures

**Figure S1.**
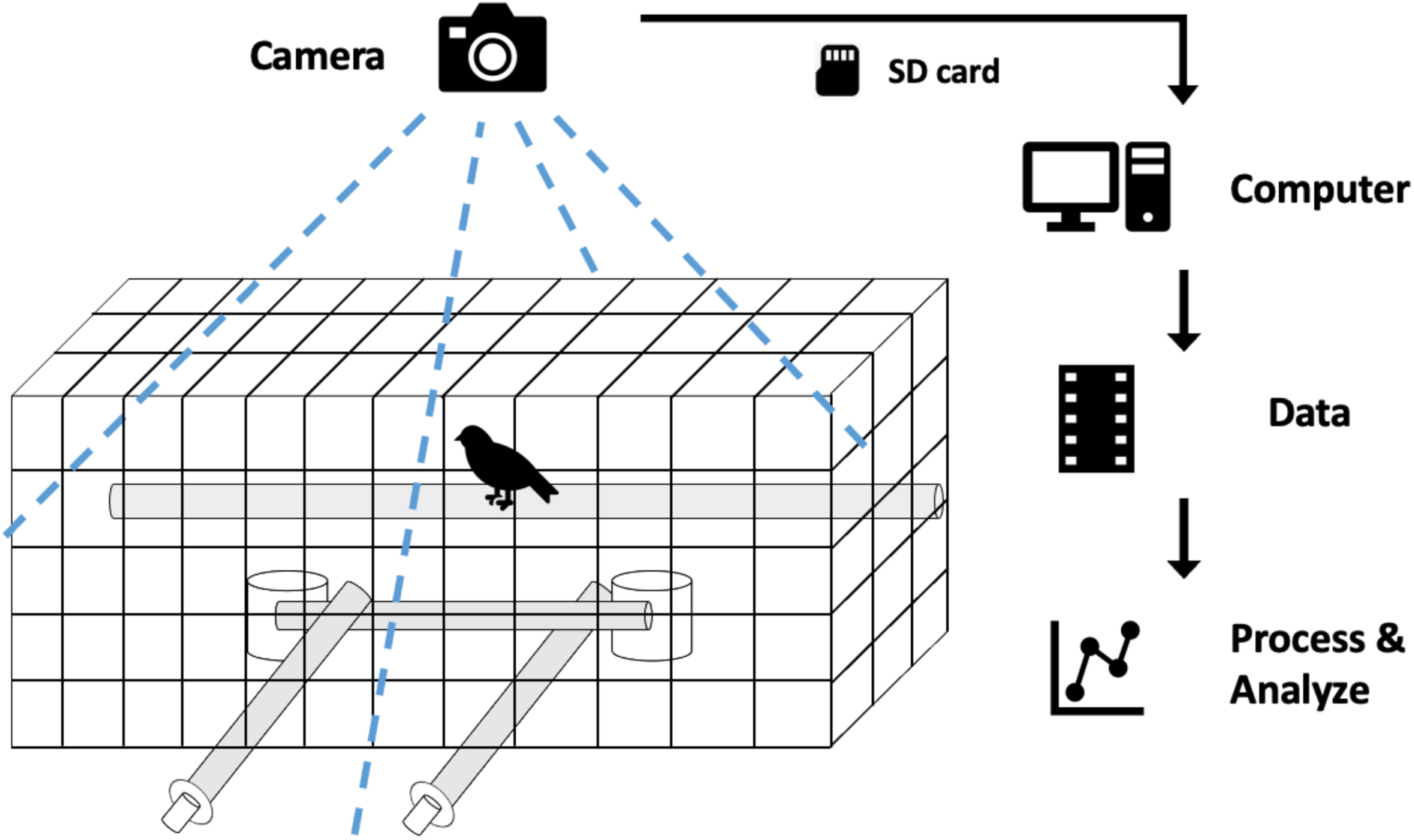
Depiction of data collection procedure.

**Figure S2.**
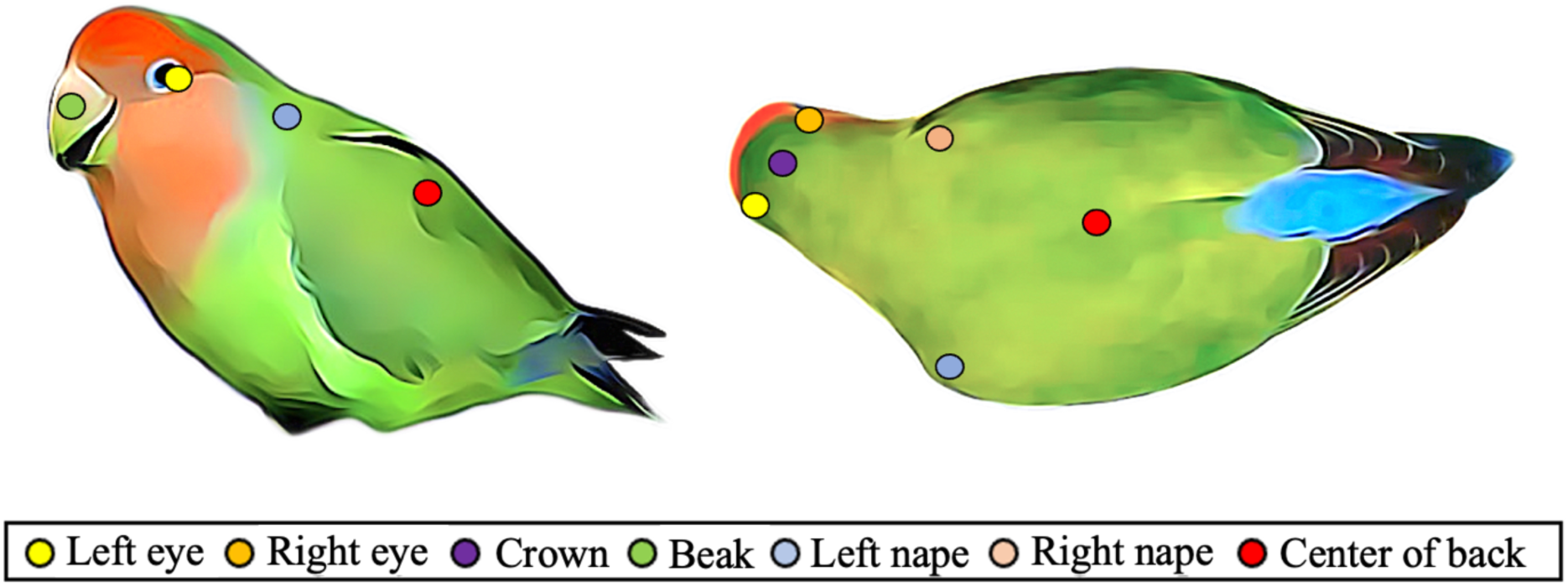
Points of interest labelled for the seven defined body parts used for training and analyses in DeepLabCut and SimBA.

**Figure S3.**
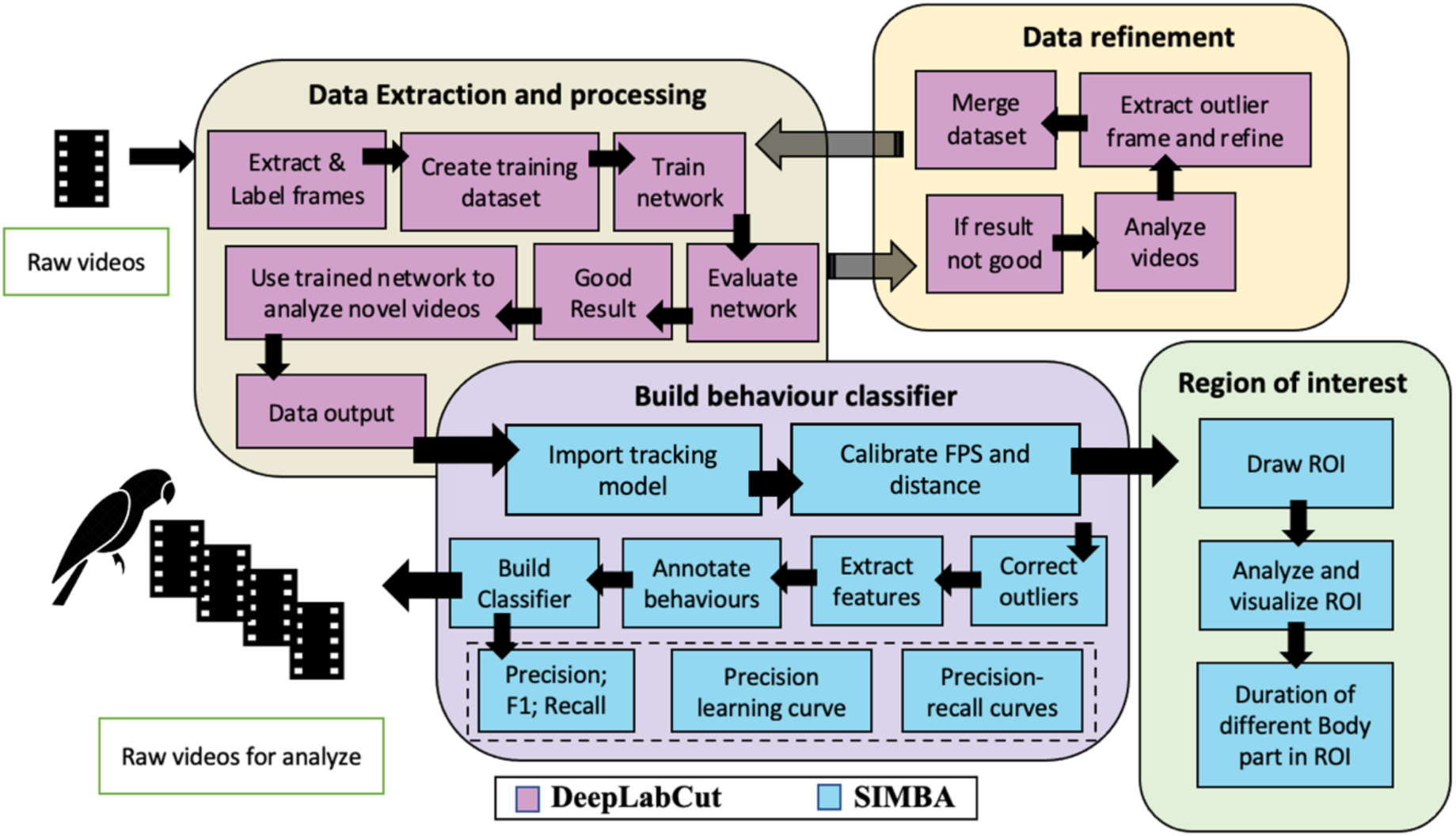
Workflow showing the generation of the pose-estimation model in DeepLabCut and the behaviour classifier in SimBA.

**Figure S4.**
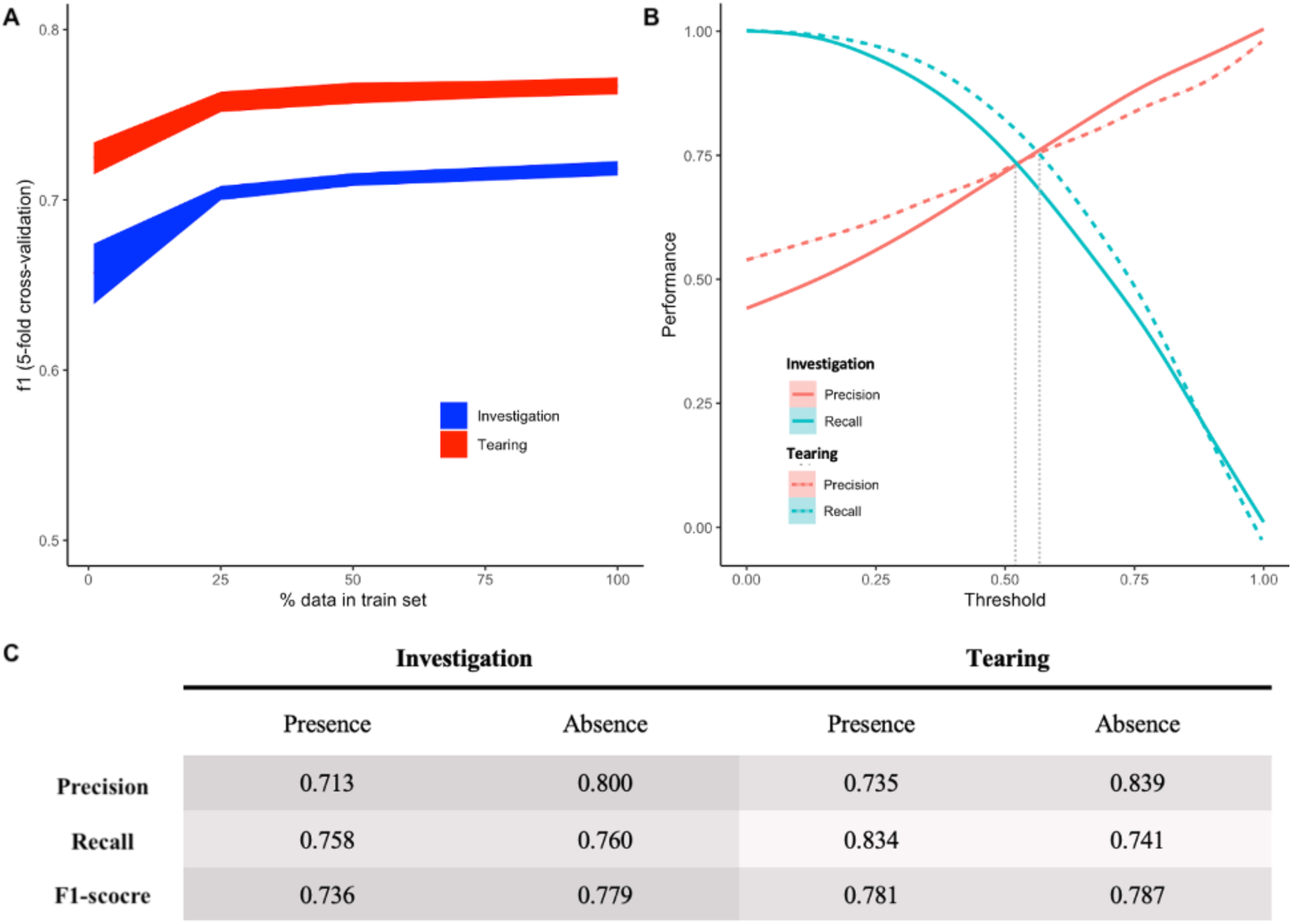
Evaluations of bird behaviour predictive classifiers. (A) Learning curves were created using 2k trees, 5 data splits (1-100%), and shuffled 5-fold cross-validation at each data split. Errors represent ± SEM. (B) Classification precision, recall, and F1 scores at different discrimination thresholds. The grey dotted line represents the discrimination threshold at the maximal F1 score. C) Mean classifier precision, recall, and F1 score evaluated by shuffled 5-fold shuffle cross-validation.

